# Physical mechanisms of nanoparticle intracellular release through membrane pore nucleation and expansion

**DOI:** 10.64898/2026.09.09.750319

**Authors:** Massimiliano Paesani, Ioana M. Ilie

## Abstract

Successful intracellular delivery requires a nanocarrier to adhere to the cell membrane, be wrapped and internalized and finally released into the cytoplasm before it is degraded. While the initial steps have been intensively studied, the intracellular release of a nanocarrier has received comparatively less attention and remains poorly understood. To understand the physical mechanisms that govern intracellular release, we investigate the escape of nanocarriers (radii 4.5*σ* and 7*σ*) from a lipid vesicle using coarse-grained simulations. We represent a nanocarrier using the metaparticle model and the membrane using the three-beads Cooke–Deserno model. Starting from a fully endocytosed state, we systematically vary the repulsion between the nanoparticle and the lipid head groups and compare three release scenarios: *passive* release, localized *inside-out* activity from a drug-like source confined within the particle and activity distributed over the entire nanoparticle. An elastic-energy analysis shows that in a tensionless membrane escape requires crossing a large pore-expansion barrier of ≈ 68*k*_*B*_*T* for the smaller particle and ≈ 110*k*_*B*_*T* for the larger, making passive release thermodynamically improbable. Our results show that the smaller particle nonetheless escapes above a threshold repulsion, whereas the larger one remains trapped in a metastable state in which local membrane pores form but fail to grow beyond the particle diameter to allow successful release. Adding activity lowers the nucleation barrier and promotes earlier pore opening, but its effectiveness depends on how it is applied, *i*.*e*., a localized internal source assists release of the smaller particle yet is insufficient to systematically free the larger one, while activity distributed across the whole nanoparticle drives complete release even in the larger case. These results show that productive escape is governed by whether the resulting pore can evolve into a geometrically accessible release pathway, providing physical design principles for intracellular delivery by nanocarriers.

## I. INTRODUCTION

The uptake of a nanoparticle by a cell can be described as a wrapping transition, in which nanoparticle-membrane adhesion drives the bilayer to bend around it, competing against the membrane bending rigidity and tension that resist deformation, until the nanocarrier is fully engulfed and internalized^1–3^. Releasing the cargo inside the cell requires the reverse, *i*.*e*., the fully wrapped, internalized nanoparticle must be released from the enclosing membrane and reach the cytosol before it is trafficked to degradative compartments^4,5^.

Intracellular release of internalized nanocarriers is both a physics and a chemistry problem. From a physical perspective, release is a rare event because the fully wrapped state is thermodynamically and mechanically stable. Essentially, adhesion to the bilayer favors retention, while the bending rigidity and tension that first drove engulfment now oppose dewrapping^6,7^. Hence, nanoparticle release requires more than membrane de-wrapping and involves breaching of the surrounding vesicle^8^. The creation of such a perturbation in the membrane requires overcoming of free-energy barriers, arising from the balance between membrane bending rigidity and the energetic penalty of exposing a free membrane edge^9–14^. In practice, membrane disruption often takes the form of a transient aqueous pore. From a chemical perspective, the composition of the nanoparticle, its surface functionalization and the surrounding environment can lower free-energy barrier of pore formation by tuning the adhesion strength^15^, the membrane’s spontaneous curvature and the line tension that resists pore formation^1,10^. Although biological and engineered escape strategies are chemically diverse, many converge on the local destabilization of the bilayer into a transient discontinuity through which the nanoparticle can escape^4,16–21^. Ionizable and fusogenic lipids (lipid mixing)^22,23^, the proton-sponge effect and pH-responsive components (osmotic and carrier swelling),^24,25^ and pore-forming peptides^26^ can thus all be understood as chemically distinct ways of driving the same physical process, *i*.*e*., lowering the energetic cost of pore opening and nanoparticle escape.

For instance, lipid nanoparticles consisting of ionizable lipids are formulated to be neutral at physiological pH but to become cationic under acidic conditions (inside the cell), where they reorganize, mix with anionic endosomal lipids and promote high-curvature structures contributing to the nanocarrier release into the cytosol^27^. The interconnected chemistry of the lipid headgroups, the shape of the lipids and their ionization properties can be tuned to stimulate distinct responses in the biological environment and thereby enhance nanocarrier delivery efficiency^17,18,27^. Furthermore, breaching the enclosing membrane is necessary but not sufficient for release, as only a small fraction of cargo escapes even from vesicles that show clear membrane damage, and some damaged vesicles release no detectable cargo^28^. As such, whether a perturbed membrane releases its content also depends on whether that pore or discontinuity can overcome additional free-energy barriers to evolve into a geometrically accessible exit pathway for the nanoparticle^29^.

From an experimental perspective, intracellular release can be assessed qualitatively, using membrane-damage or leakage probes, and quantitatively, using cytosolic-access assays^30,31^. The bottleneck arises because the pore that allows a nanoparticle to escape is small and transient, its formation governed by a nucleation barrier and by the edge tension of the pore^9,10,14,32^. With nanoparticle escape being a rare and fast event, imaging the process directly is difficult, and fluorescence-based readouts of cargo localization are practical approaches^33,34^. However, interpreting them remains challenging, as the strong punctate signal from the cargo still trapped in vesicles dominates over the much weaker diffuse cytosolic signal that marks successful release^4,16,18,30^. Even when quantification becomes possible, reported escape fractions are often only a few percent^33^ with much of the internalized dose recycled back from the cell^34^.

Computer simulations are complementary to experiment providing insight into the details of membrane deformation, pore nucleation and into the physical mechanisms that govern nanoparticle uptake^35**?**–42^ and release^8^. In particular, coarse-grained models calibrated on experimental elastic properties^43–45^ have explored the effect of adhesion and membrane bending in the cellular uptake of nanoparticles^1,3,46–51^. Additionally, they provided insight into the resistance of lipid bilayers under deformation and rupture under tension and how a nucleation barrier and the line tension govern the opening of a pore in a bilayer^9,10^. Furthermore, the escape of nanoparticles internalized via passive endocytosis was reproduced by lowering the interaction with the bilayer^8^. Small nanocarriers were successfully released, yet larger ones remained trapped unless the membrane was placed under tension, achieved through particle expansion^8^. In this context, release is an equilibrium outcome, driven by reduced adhesion and stored elastic stress. The intracellular environment, however, is not at equilibrium as it sustains active, energy-consuming processes that continuously do work on enclosed cargo. Whether such activity can breach the confining bilayer, and how this depends on particle size, confinement and membrane elasticity, remains open.

Here we investigate the physical mechanisms of nanoparticle release under passive and active conditions using coarsegrained simulations. Starting from a quasi-spherical nanoparticle fully engulfed in a lipid vesicle, we first increase its repulsion to the membrane to drive passive release and then introduce activity to probe the destabilization of the enclosing vesicle. Our results show that, under passive conditions, the smaller nanoparticle is released once its repulsion to the membrane is strong enough to stretch the enclosing bilayer past the point of pore nucleation, whereas a larger nanoparticle of the same shape remains trapped in a metastable, partially opened vesicle. Activity lowers the effective barrier by promoting earlier pore opening and enhanced release, but its efficiency depends on how the perturbation is applied. A localized active source inside the carrier facilitates release of the small nanoparticle, yet still is insufficient to enhance the escape of the larger one. Activity distributed over the whole carrier produces a more coordinated stress on the vesicle and enables complete release even for large nanoparticles. Nanoparticle escape is thus governed by a sequence of free-energy barriers, from pore nucleation to expansion beyond the particle diameter, that must be overcome to open a geometrically accessible pathway for translocation.

## II. SIMULATIONS DETAILS

We investigate the intracellular release of an endocytosed nanoparticle from a lipid vesicle using coarse-grained simulations. A nanoparticle is represented using the recently introduced metaparticle (MP) model^52^ and the membrane using the Cooke–Deserno three-bead lipid model^43,44^. Briefly, in the metaparticle model, a particle is represented by a collection of beads placed in symmetric fashion at the vertices of a polyhedral scaffold and interconnected by spring-like potentials^52^. Here we used MP_60_ and MP_180_ as model systems (Fig. S1), consisting of 60 and 180 interconnected beads, respectively^52^. Both models share the same icosahedral symmetry but differ in size, *i*.*e*., they have diameters of 9 *σ* and 14 *σ*, respectively, with *σ* = 1nm. In the Cooke–Deserno model, each lipid is coarse-grained into one head bead and two tail beads linked by nearly inextensible bonds and interacting via generic pair potentials^43^. The details of both models are in the Supplementary Information. The interactions between the metaparticle and membrane beads are modeled using a modified 12–6 Lennard–Jones potential with independently tunable repulsive and attractive contributions (Eq. S1)^46^. All simulations were performed in LAMMPS in reduced Lennard–Jones units, with *ε* and *σ* defining the units of energy and length, respectively, and the time unit 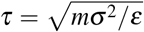. The dynamics of the system was propagated using a velocity-Verlet integrator with the temperature maintained at *T* = 1.1*ε/k*_*B*_ by means of a Langevin thermostat.

A typical simulation is started from a fully endocytosed state in which a nanocarrier is encapsulated in a closed lipid vesicle (Fig. 1(a)). The encapsulated configuration was obtained from a successful endocytic event reported in our previous work^46^, in which the nanoparticle was fully wrapped by the membrane and enclosed within a lipid vesicle. To use this state as the starting point for the release simulations, we extracted the vesicle containing the encapsulated nanocarrier from the final uptake configuration. The remaining membrane region, which was no longer directly associated with the vesicle or the nanocarrier, was removed. The lipid vesicle and the enclosed nanoparticle (≈ 1150 lipids and ≈ 2140 lipids for MP_60_ and MP_180_, respectively) were then used as the starting conformation for the release simulations. The reduced system was placed in a cubic box with periodic boundary conditions and edge length 43.2 *σ*, preserving the bead number density of the generated initial system^46^. Each system was run for 1.0 · 10^7^ steps, with a timestep *δt* = 0.001 *τ* for a total simulated time 1.0 · 10^4^ *τ*, with conformations saved every 500 timesteps. To ensure statistical significance, five independent copies of each system were simulated under the same conditions using different random seeds.

**FIG 1.**
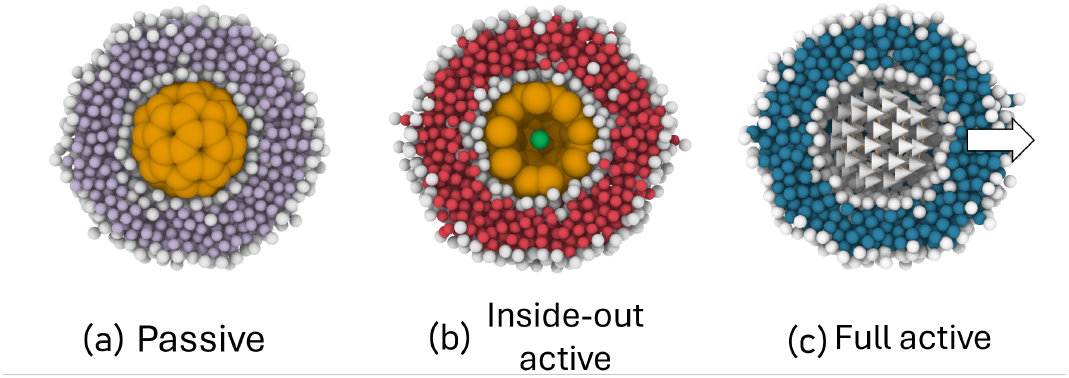
Schematic representation of the three release systems considered in this work: (a) *passive*, where release is controlled by tuning the repulsive interaction between the nanoparticle (orange) and the lipid head beads (heads shown in white and tails in light purple); (b) *inside-out active*, where an active drug-like particle (green bead) is confined inside the nanocarrier and generates an internal driving force for release, and (c) *full active*, where all beads of the nanoparticle are active and collectively drive the release process.

## III. RESULTS

To investigate the mechanisms of nanoparticle escape, we studied three distinct systems starting from a nanoparticle encapsulated in a lipid vesicle (Fig. 1). First, we investigated the conditions under which a nanoparticle by thermal fluctuations alone is passively released from the bilayer vesicle (referred to as *passive* system). As such, we varied the repulsive interaction between the nanoparticle and the lipid head beads, *ε*_rep_ in the 1-9*ε* interval. Second, to mimic a changing physicochemical environment between the endosomal interior and the cytosol, we placed a self-propelled particle inside the nanoparticle (referred to as *inside-out-active*). For this, we applied a self-propulsion force along the instantaneous velocity vector of magnitude 5 *ε/σ* on the internal particle, thereby exerting an internal mechanical perturbation on the nanoparticle and, indirectly, on the surrounding vesicle (green particle in Fig. 1(b)). Third, to mimic a coordinated external driving force, we applied an active force of 0.1*ε/σ* to every bead of the metaparticle, all directed along the x axis (referred to as *full active*).

### A. Passive release

To understand nanocarrier release relative to particle size, we considered two nanoparticles of distinct sizes with icosahedral symmetry, *i*.*e*., MP_60_ and MP_180_ (diameter ≈ 9*σ* and 14*σ*, respectively), subjected to thermal fluctuations under *passive* conditions. A typical system is initialized starting from the fully encapsulated state, with a nanoparticle enclosed in a lipid bilayer (Fig. 1(a)), extracted from our previous work on cellular uptake of nanoparticles^46^. The resulting vesicles differ in lipid content and size, containing approximately 1150 lipids for MP_60_ (outer vesicle diameter ≈ 20*σ*) and about 2140 for MP_180_ (outer vesicle diameter ≈ 25*σ*). To enhance the probability of escape, we fix the attraction between the metaparticle and lipid beads at *ε*_attr_ = 1*ε* and systematically vary the repulsive interactions, *ε*_rep_, in the 1-9*ε* range. For each condition, we perform five independent simulations to ensure statistical robustness. Our results show that with increasing *ε*_rep_, three distinct regimes emerge. At low repulsion, the nanoparticles remain fully encapsulated in the lipid vesicle over the timescale of the simulations. At intermediate repulsion, a metastable state emerges, in which pores form in the bilayer without complete nanoparticle escape on the simulation timescales. At strong repulsion, MP_60_ is released (Fig. 2(a) and Fig. 3(a)). The larger MP_180_ remains encapsulated in the vesicle independently of the repulsion strength.

**FIG 2.**
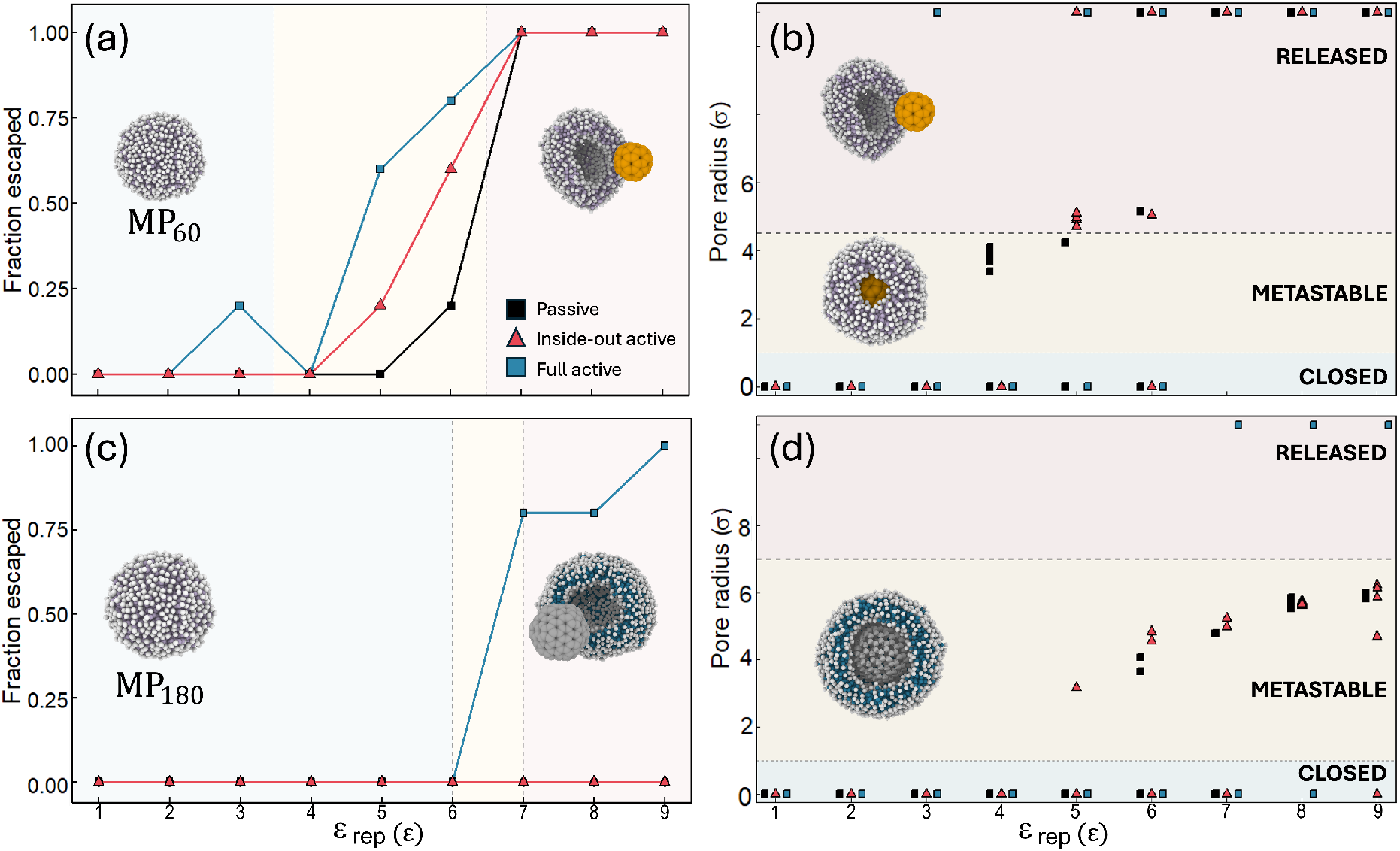
Fraction of successfully escaped nanoparticles and characteristic pore radius as a function of the MP-head-bead repulsion strength. Panels (a) and (c) show the fraction of released particles for MP_60_ and MP_180_, respectively, calculated over five independent simulations. Panels (b) and (d) show the average emerging metastable pore radius for each replicate. The pore radius is calculated by projecting the pore-forming outer-leaflet head-beads onto a target plane and measuring the distance between the exit axis and the nearest projected head bead after subtracting the effective head-bead dimension. For releasing trajectories, the symbols only highlight the fact that the nanocarrier escapes the vesicle. For the metastable states, the symbols show the average radius of the metastable pore. Highlighted are the passive (black squares), *inside-out* active (red triangles) and fully active systems (blue squares).

**FIG 3.**
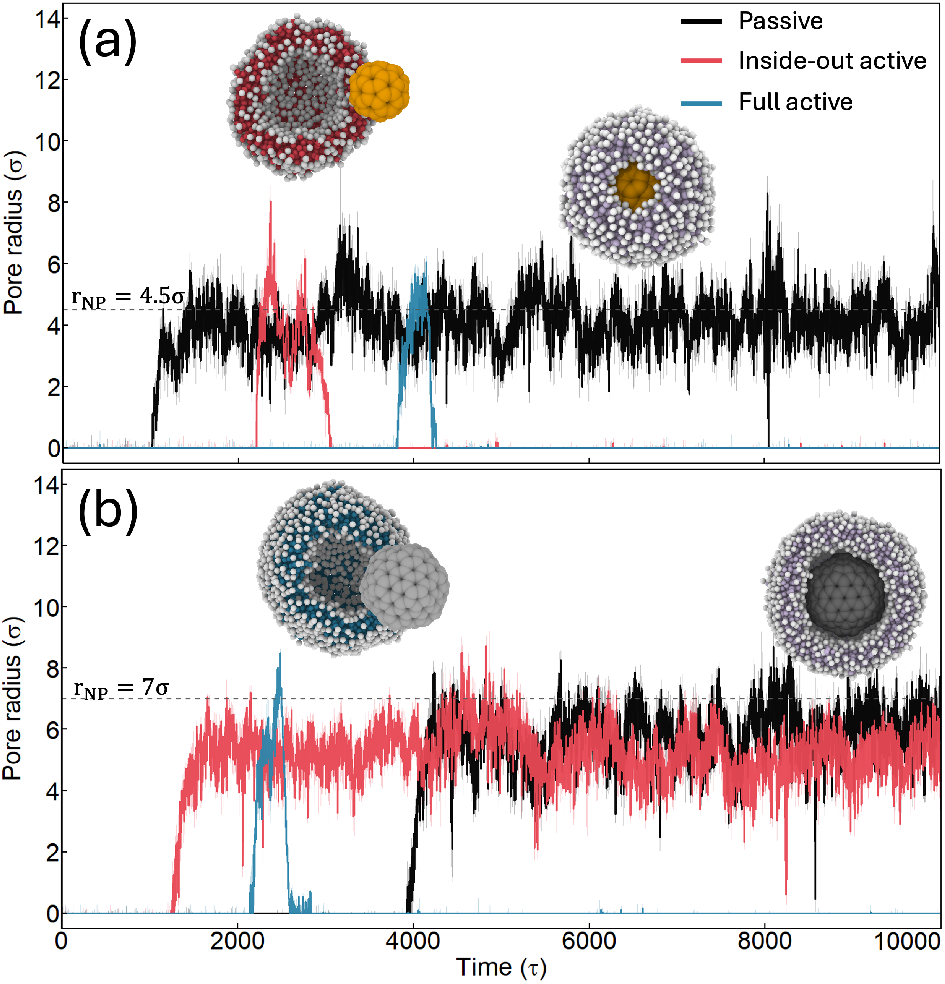
Pore opening dynamics. Representative pore-radius trajectories for passive, inside-out active and fully active systems showing metastable and release-competent states. (a) For MP_60_, the selected trajectories correspond to *ε*_rep_ = 4*ε*, 9*ε*, and 6*ε* for the passive (black lines), inside-out active (red lines) and fully active systems (blue lines), respectively. (b) For MP_180_, the corresponding trajectories are shown at *ε*_rep_ = 8*ε*, 7*ε*, and 8*ε*, respectively. Highlighted are the running medians over 5 frames. The horizontal dashed lines indicate the nanoparticle radii, *r*_NP_ = 4.5*σ* for MP_60_ and *r*_NP_ = 7*σ* for MP_180_. For MP_60_, a pore opens around 1000 *τ* and remains in this metastable state over the entire simulation (black line, passive case). The active trajectories (red blue lines) show pore opening followed by closing after successful nanoparticle release. For MP_180_, the passive and inside-out active trajectories develop persistent pores fluctuating in size though remaining below the nanoparticle radius, whereas full activation drives pore expansion beyond the nanoparticle radius, release and pore closing.

At low repulsion values, *ε*_rep_ *<* 4*ε*, the nanoparticles remain fully embedded within the lipid vesicle independently of their size (black points in Fig. 2(a-d)). In this regime, the perturbation generated by the thermal motion is not sufficient to induce a strong local lipid organization around the nanoparticle and induce tension in the membrane to allow release. As a result, the encapsulating vesicle is stable and no pore is formed in the membrane (black lines in Figs 2(b) and (d)), preserving the system in a fully enclosed state.

For intermediate values of *ε*_rep_ ∈ 4 − 6*ε*, the passive MP_60_ locally perturbs the bilayer resulting in an increased fraction of nanoparticles released from the vesicle (Fig. 2(a)), consistent with the emergence of pores in the lipid vesicle (Fig. 2(b)). This behavior is observed in four replicas at *ε*_rep_ = 4*ε*, one of five replicates at *ε*_rep_ = 5*ε* and in two replicates at 6*ε* (black profile in Fig. 3(a,b)). For *ε*_rep_ = 4*ε*, a metastable pore of radius *r*_*p*_ ≈ 4*σ* emerges, below the particle radius (black line in Fig. 3(a)). Despite excursions to radii above the particle radius, these nanoparticles do not escape, as release requires the pore to sustain an enlargement beyond the nanoparticle radius for the nanocarrier to be able to diffuse out of it. Mechanistically, to initiate release, the nanocarrier induces local rearrangements in the lipid bilayer, which lead to a progressive membrane destabilization and pore formation. The layer of head beads facing the nanocarrier rearranges and progressively merges with the outer head-bead layer, leading to the formation of a pore (Movie S1). Increasing the repulsion widens the metastable pore, as its radius grows from ≈ 3 −4*σ* at *ε*_rep_ = 4*ε* to ≈ 5*σ* at 5-6*ε* (Fig. 2, (Fig. S3). This metastable pore is stable over the simulation timescales and only in an isolated case at *ε*_rep_ = 6*ε* progresses towards further enlargement or nanoparticle release. For MP_180_ increasing the repulsion towards the lipid heads within the same ranges, does not result in any nanoparticle release scenarios (Fig. 2(c), Fig. 3(b)). However, similar to MP_60_ it leads to the emergence of metastable pores of ≈ 4*σ* in pore in the lipid vesicle at *ε*_rep_ = 6*ε* (Fig. 2(d)).

For *ε*_rep_ *>* 6*ε*, the pore in the resulting vesicle expands beyond the nanoparticle radius for an extended time (Fig. S3) for MP_60_, which is geometrically large enough to allow nanocarrier release. In this state, thermal fluctuations and the collisions with the lipid beads drive the system out of the metastable state, promoting pore expansion and nanoparticle release. Interestingly, after release the pore reseals and the vesicle closes off (Fig. S3). For the larger MP_180_, increasing *ε*_rep_ leads to membrane rearrangement and pore enlargement from ≈ 5*σ* at *ε*_rep_ = 7*ε* to ≈ 6*σ* at *ε*_rep_ = 9*ε* (Fig. 2(d)). However, this pore is not large enough to complete translocation, leaving the nanocarrier still trapped in a metastable state, suggesting that, for MP_180_, the thermodynamic barrier associated with further pore expansion remains too high to allow release.

### B. Activity-mediated intracellular release

To probe this hypothesis and enhance release probability, we introduced activity into the system via two distinct mechanisms. First, we mimicked an internally generated mechanical perturbation, in which active motion originating from inside the nanocarrier propagates to the surrounding vesicle. In this case, activity arises from a particle confined within the nanocarrier (green particle in Fig. 1(b)) and self-propelled with a force of 5*ε/σ* along its instantaneous velocity vector, further referred to as *inside-out* activation. This setup is inspired by biological escape scenarios where protonation, swelling, osmotic stress or rupture-like events build up within the endosomal compartment or the carrier itself and locally promote release^17,25,30,53^. Second, we modeled a more collective, membrane-active mechanism, analogous to ionizable carriers whose components cooperatively destabilize the endosomal membrane upon acidification rather than acting through a single localized perturbation^17,54,55^. Here, activity is homogeneously distributed over all nanocarrier beads, each of which experiences a constant force of magnitude 0.1*ε/σ* along the *x* direction.

Our results show that, for MP_60_, activation enhances the regimes identified in the passive systems by driving the system out of equilibrium. For *ε*_rep_ *<* 5*ε*, the nanocarrier remains encapsulated in the closed vesicle, indicating that the induced perturbation is still insufficient to produce sustained membrane remodeling and its associated rearrangement (red profiles in Fig. 2(a) and Fig. S3). In the fully active system, nanoparticle release is observed in one replica at *ε*_rep_ = 3*ε* (Fig. 2(a)), suggesting that mechanically induced stress can create defects in the membrane that destabilize the vesicle.

For higher repulsion, activity shifts the system behavior and the intermediate regime becomes narrower. At *ε*_rep_ = 5*ε*, complete release is observed in one of five MP_60_ simulations, while in the remaining four, metastable pores form with *r*_*p*_ ≈ 5*σ* (Fig. 2(a)). In the successfully released case, the pore grows to radii significantly larger than the nanoparticle radius (up to ≈ 9*σ*), the nanocarrier leaves the vesicle, which subsequently closes (red line in Fig. 3(a)). In the metastable systems, despite the formation of a release competent pore, these trajectories do not develop a persistent pathway for complete release with MP_60_ remaining confined in the half open vesicle. This does not come as a surprise, as the nanocarriers must diffuse out of the vesicle, which may occur on longer timescales. Under fully active conditions, MP_60_ successfully escapes the vesicle in three of the five runs at *ε*_rep_ = 6*ε* (Fig. 2(a), Fig. S3). Beyond the 7*ε* repulsion threshold, MP_60_ is released in all runs from the vesicle independently of the type of activity. For MP_180_, activity has a pronounced effect for repulsion strengths *ε*_rep_ ≥ 7*ε*. Similarly as for the passive case, the inside-out active systems develop metastable pores with radii ranging from ≈ 5*σ* for *ε*_rep_ = 6*ε* to ≈ 6*σ* for *ε*_rep_ = 9*ε*. Nevertheless, MP_180_ remains encapsulated in the vesicle in all runs sampling the *inside-out* activation (red line in Fig. 3(b)). Full activation on the other hand, results in successful escape in four of five runs at *ε*_rep_ = 7*ε* and 8*ε*, and in all five runs at *ε*_rep_ = 9*ε* (blue line in Fig. 3(b), Fig. S4).

### C. Release mechanisms

Pore formation is a nucleation process, that arises from the competition between the energy cost of creating an exposed pore edge and the energy gain from relieving membrane stress^8,10,14,56^. Hence, the membrane needs to overcome a free-energy barrier, associated with the emergence of local defects to create a pore. Overcoming this barrier is a stochastic event and therefore pore formation can occur on different timescales under identical conditions, as observed throughout the *passive* and the *inside-out* systems. To understand the release mechanisms, we focus on the systems with successful release events starting from the early initiation of the pore.

For both MP_60_ and MP_180_, a successful release is initiated by the collisions between the metaparticle beads and the lipid head groups, which destabilize the membrane and locally rearrange the membrane beads (first snapshot in Fig. 4(a)). These interactions thereby increase bilayer frustration and favor the formation of local defects. As the membrane responds, the lipid head groups move away from the most stressed region, the two leaflets become locally asymmetric, and the lipid tails become more exposed and disordered near the thinning site (second snapshot in Fig. 4(a)). This creates an intermediate state, in which hydrophobic tails are transiently exposed to solvent, consistent with previous simulations^9,10,14,57^. The defect then evolves from a hydrophobic pore precursor into a small hydrophilic pore as head groups from the inner and outer leaflets rearrange to line the pore rim (second snapshot in Fig. 4(a)). This transition stabilizes the pore edge, relieves elastic stress and is accompanied by an increase in vesicle outer radius (dark orange profile in Fig. 4(b)). Subsequently, as the pore expands beyond the diameter of the nanocarrier, the nanoparticle is able to escape through the opening (third snapshot in Fig. 4(a)) and the pore closes (fourth snapshot in Fig. 4(a)). While this behavior is generic for the tested nanoparticles and is independent of whether the system is passive or inside-out activated, the fully active nanocarrier can also follow a different release pathway. Specifically, due to its stronger activity, it can induce stronger deformations of the surrounding vesicle, giving rise to local membrane perturbations that are not always localized with the nanocarrier displacement direction. Consequently, pore opening may occur at a remote location rather than along the direction of applied activity, enabling the nanocarrier to escape by rolling out of the vesicle (Fig. S5).

**FIG 4.**
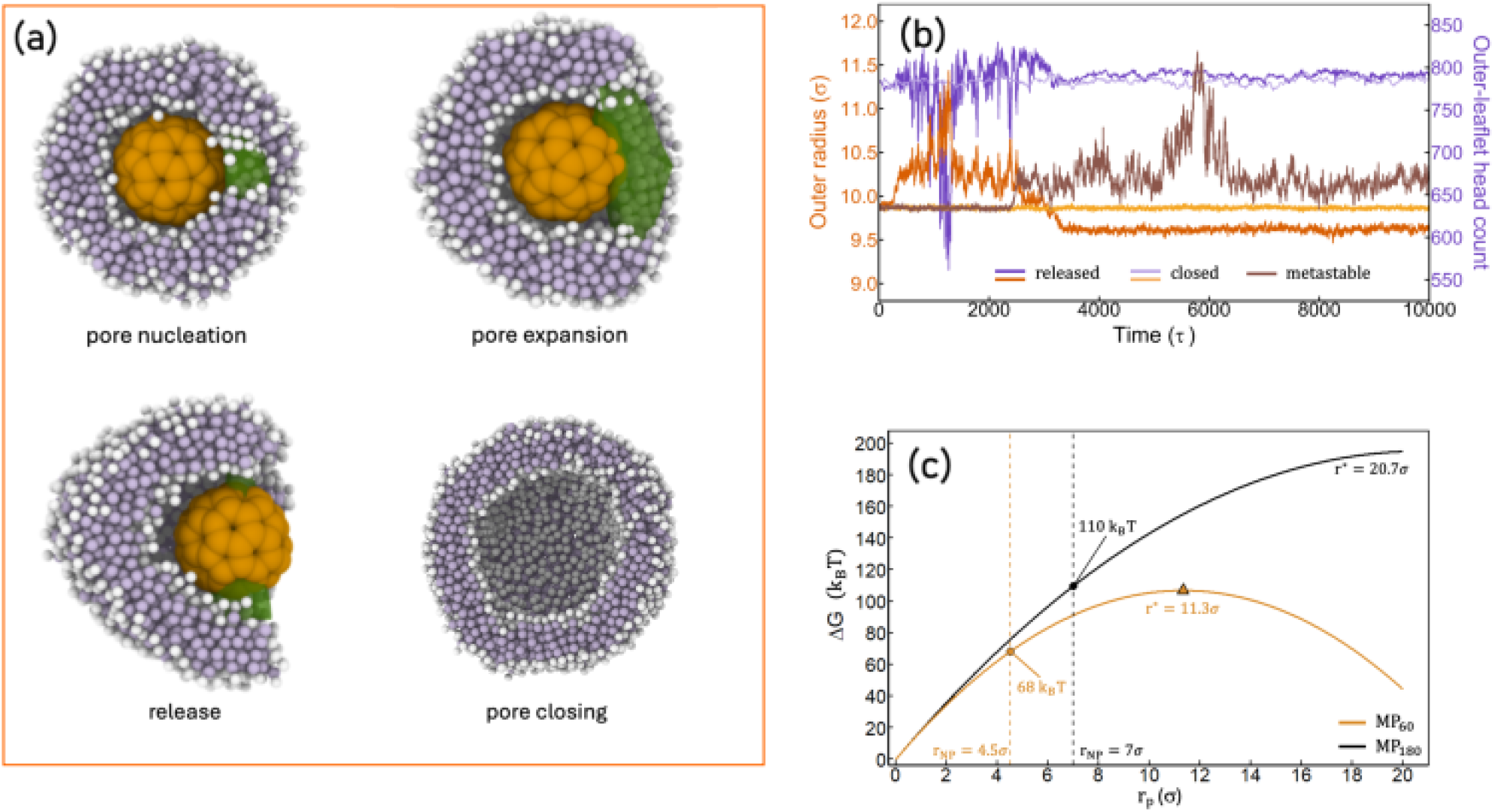
Mechanism and free-energy landscape of nanoparticle release from a lipid vesicle. (a) Representative sequence of membrane destabilization and pore nucleation, pore expansion, nanoparticle release and pore closing (sliced view). (b) Time evolution of the outer vesicle radius and outer-leaflet head-bead count for passive MP_60_ at *ε*_rep_ = 5*ε* (light orange and light purple lines, respectively), for which the vesicle remains closed and at *ε*_rep_ = 7*ε*, for which membrane rearrangement precedes pore opening and nanoparticle release (dark orange and dark purple lines). The brown profile highlights a metasble state at *ε*_rep_ = 5*ε inside out* active. (c) Free-energy cost of forming a pore of radius *r*_*p*_ in tensionless vesicles enclosing MP_60_ (orange) and MP_180_ (black), calculated from Eq. S2 using *γ* = 3 *k*_*B*_*T/σ, κ* = 13 *k*_*B*_*T*, and *κ*_*G*_ = −11.5 *k*_*B*_*T*. The vesicle mid-plane radii are *R*_0_ = 7.4*σ* and 10*σ* for MP_60_ and MP_180_, respectively. The dashed vertical lines indicate pore radii equal to the corresponding MP radii, *r*_*p*_ = 4.5 *σ* and 7.0 *σ*, for which the free-energy costs are approximately 68 *k*_*B*_*T* and 110 *k*_*B*_*T*. The critical radii separating pore closing from continued expansion are *r*^∗^ = 11.3*σ* for MP_60_ and *r*^∗^ = 20.7*σ* for MP_180_.

For successful release to occur, the system must overcome a series of free-energy barriers, from the nucleation of a defect to the opening of the pore and its expansion to the size of the nanoparticle. Their combined effect determines whether the vesicle merely swells, forms metastable pores or releases its cargo. In the passive case, at moderate repulsion, the vesicle around MP_60_ is assumed tensionless and its outer radius equilibrates at 9.86 *σ* (light orange line in Fig. 4(b)). Releasing a nanoparticle from a tensionless vesicle requires the formation and the enlargement of the pore to a radius comparable to or larger than the nanoparticle radius, *i*.*e*., 4.5*σ* for MP_60_. Following elasticity theory, the barrier for opening a releasecompetent pore in a tensionless vesicle is given by 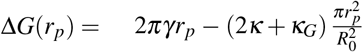, with *γ* the line tension of the pore edge, *κ* and *κ*_*G*_ the bending and the Gaussian bending rigidities, respectively, *K*_*S*_ is the stretching modulus of the membrane, *R*_0_ the membrane mid-plane radius of the surrounding vesicle and *r*_*p*_ the pore radius^8^. Here we chose *κ* = 13*k*_*B*_*T, κ*_*G*_ = −11.5*k*_*B*_*T, γ* = 3*k*_*B*_*T/σ* as measured from the original Cooke-Deserno model^43,58^ and *K*_*S*_ = 250*mJ/m*^210^. MP_60_ has a radius of 4.5*σ* and is initially wrapped by a vesicle of mid-membrane radius 7.4*σ*. Hence, the free-energy barrier that the system needs to overcome to nucleate a release-competent pore is of about 68*k*_*B*_*T* (Fig. 4(c), Eq. S2), which can only rarely be overcome by thermal fluctuations, consistent with the non-releasing states. As repulsion towards the lipid heads is increased, the nanoparticle collides more strongly with the inner leaflet. The intensified collisions build up an internal pressure that drives a rearrangement of the lipids and the emergence of a transient swelling of the vesicle (dark orange line in Fig. 4(b)).

For instance, at a repulsion of *ε*_rep_ = 7*ε*, in the early stages (first 100 *τ*), while the membrane is still closed, the swelling stretches the bilayer as the outer vesicle radius increases to about 10.2*σ*, with the outer leaflet lipid count fluctuating around 780 (Fig. S6), corresponding to an area strain of 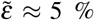 (Eq. S4). This strain imposes a transient tension 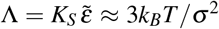, close to the rupture threshold of the bilayer, which lowers the nucleation barrier for a pore, Δ*G* = *πγ*^2^*/*Λ ≈ 9 *k*_*B*_*T* (Eq. S6). Upon activation, the net effect is essentially the same. For example, using the *inside-out* mechanism at *ε*_rep_ = 5*ε*, the vesicle swells at an equilibrated outer radius of about 9.9*σ*, comparable to the passive case. As the internal drive builds up, the vesicle swells while still closed, its outer radius reaching ≈ 10.2*σ* with the outer-leaflet lipid count constant. Importantly, in most *passive* and *insideout active* systems, the pores reseal after nanoparticle release, which is evident from the reduced outer radius of the vesicle, which equilibrates at 9.6*σ* (dark orange line in Fig. 4(b)). This is happening because the radius for the opening remains lower than the critical radius of the tensionless exit barrier^59^ 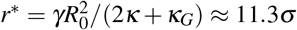 (Eq. S3). Hence, once the particle has passed, the pore lies on the resealing side of the freeenergy barrier and it is thermodynamically more favorable to close. In contrast, in some cases the vesicle does not always reseal. Instead, the pore keeps expanding, the outer radius grows as the membrane opens up completely, relaxing into an extended bilayer from which the particle is released. Hence, across the systems that successfully release, the pore opens under a mechanical stress that emerges from a confined particle of fixed size colliding with the inner leaflet of its surrounding vesicle, reaching the same outcome as previous simulations that opened pores by an externally applied membrane tension^9,10^, or by growing the encapsulated particle^8^.

In the metastable regime, the vesicle around MP_60_ fluctuates around an equilibrium radius of ≈ 9.86*σ* (*ε*_rep_ = 5*ε, inside out* active) after which it slightly increases and fluctuates around an equilibrium outer radius of ≈ 10.3*σ*. (brown profile in Fig. 4(b)). This increased outer radius emerges together with the formation of a pore within the first 2000 *τ*. While the outer vesicle experiences some swelling and recovering around 6000 *τ*, the pore that forms, neither reseals nor grows to release the nanoparticle, and remains stable at a pore radius of approximately the nanoparticle radius (Fig. S3). Evaluated at this swollen outer vesicle radius (mid-plane radius *R*_0_ ≈ 7.8*σ*), the tensionless release barrier for full release becomes Δ*G* ≈ 70*k*_*B*_*T* and a critical radius 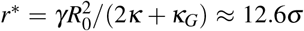. We interpret this metastable pore using an adaptation of the constant-area pore free-energy introduced for flat bilayers^9^, extended here to a curved vesicle. Hence, the total membrane free-energy when a pore of radius *r*_*p*_ is present, is given by:

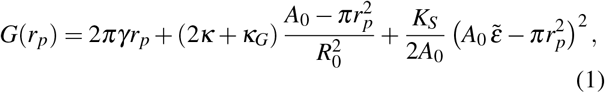

with *A*_0_ the tensionless area of the outer leaflet and 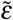 its area strain. Here, the first term on the r.h.s. represents the line-tension cost of the pore rim, the is the curvature energy of the remaining membrane, which is lowered as the pore removes curved area, and the last term represents the elastic energy stored in the residual area strain after the pore has absorbed area 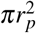. For MP_60_, Δ*G*(*r*_*p*_) allows the formation of a metastable pore within a strain window, 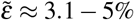, below which the pore reseals and above which any pore will grow (Figs. S3, S7). Within this window the metastable pore minimum is between *r*_*p*_ ≈ 2.4 − 4.1*σ* behind a barrier of order 10*k*_*B*_*T* (Fig. S7), approaching the nanoparticle radius of 4.5*σ* at the upper end of the window. Indeed, in some cases the pore radius equilibrated at values slightly higher than the nanoparticle radius (Fig. 2(b)) and we ascribe this partly to the finite width of the pore rim and partly to the finite duration of our simulations.

the second is the curvature energy of the remaining membrane, which is lowered as the pore removes curved area,

## IV. DISCUSSION AND CONCLUSION

Nanoparticles are predominantly taken up by cells through the endocytic pathway, in which they become enclosed within membrane-bound intracellular vesicles. Escape from these endocytic compartments is a critical and often a rate-limiting step in effective intracellular delivery. Several mechanisms have been proposed to facilitate the escape. These include membrane destabilization, in which direct interactions between the nanoparticle surface and the endosomal lipid bilayer increase membrane permeability and promote pore formation^4,18,30^.

Physically and chemically, breaching the enclosing membrane is necessary but not sufficient for release as even vesicles with clear membrane damage release only a small fraction of their cargo and some show no detectable release at all^28^. Our results show that one release strategy lies in the formation of a defect taking the shape of a pore that can grow beyond the nanocarrier diameter to allow nanoparticle escape. For this, we used a minimal coarse-grained model to understand the physical mechanisms by which nanocarrier size, confinement and activity drive vesicle destabilization, pore formation and nanoparticle escape. To this end, we compare three different systems: a *passive* system, in which the dynamics of the nanoparticle is unperturbed, an *inside-out active* system, in which changes of the intracellular environment are mimicked by placing a self-propelled particle inside the nanocarrier, and a *full active* system, in which each bead of the nanocarrier is driven by a constant force along a fixed direction.

Our results are three-fold. First, under passive conditions, the release of the nanoparticle is governed by its size. Small nanoparticles (diameter ≈ 9 nm) are released from the surrounding vesicle by simply increasing the repulsion between the nanoparticle and the lipid head beads. In this case, the enhanced repulsion locally perturbs the lipid bilayer, gradually leading to membrane thinning, defect formation and finally pore nucleation and escape. Larger nanoparticles (diameter ≈ 14 nm) fail to complete the escape under the same conditions. Although local pore openings form, they remain below the nanoparticle diameter and do not grow to allow release, consistent with experiments showing that membrane damaged vesicles fail to release their cargo^28^. Physically, pore formation is governed by the balance between the line tension, which acts to close the pore, and the energy gained by relieving membrane stress upon pore opening^9,10^. To escape, a pore of at least the size of the nanoparticle should open in the enclosing vesicle. Hence, the smaller nanocarriers require a 4.5 nm pore while the larger ones require a 7 nm pore, which increase the release barrier from ≈ 68 *k*_*B*_*T* to ≈ 110 *k*_*B*_*T*, respectively. Additionally, the smaller nanocarriers have a higher curvature relative to the membrane thickness as compared to larger ones, thereby concentrating the wrapping-induced stress over a small membrane surface area, further contributing to lowering the nucleation free-energy barrier^8^. For larger nanocarriers, the stress is distributed over a broader surface area thereby reducing the local driving effects for pore opening and growth and implicitly having a higher free-energy barrier to overcome^60^.

Second, our results inform on the enhanced release observed under active conditions. In particular, a self-propelled particle confined inside the nanoparticle experiences frequent collisions with the nanoparticle interior. This generates an internal pressure in the surrounding vesicle, which increases the mechanical tension in the bilayer, leading to an increase in vesicle volume. When this tension exceeds the membrane’s mechanical stability threshold, it can induce pore formation or widen metastable pores. For the small nanoparticles, activity enhancement can facilitate the release of the vesicle contents. For the larger nanoparticles, the same mechanism is insufficient to drive release because the collisions of the self-propelled particle with the nanoparticle wall are less frequent and generate too little internal pressure to enhance sustained pore expansion. More broadly, our results connect to previous work on active particles under confinement, where self-propulsion generates effective pressure on surrounding boundaries^61^. In confined active systems, persistent self-propulsion leads to boundary accumulation and generates a non-equilibrium mechanical pressure on the confining walls^61,62^. In our case, this wall pressure is exerted on a soft vesicle membrane, so the active stress translates into bilayer tension, enhanced membrane fluctuations and an increased likelihood of pore nucleation. Although *inside-out* activation does not change the underlying release mechanism, it influences the pore size and its stability. The self-propelled particle generates an internal active pressure within the nanocarrier that is transmitted to the outer vesicle membrane. Therefore, it induces a local mechanical stress on the membrane that contributes favorably to reducing the free-energy barrier for pore formation. This effect is insufficient for MP_180_, because its cavity is larger than that of MP_60_ and therefore the self-propelled particle diffuses more freely, collides less frequently with the carrier wall and transfers less energy to the membrane. In MP_60_, the stronger confinement increases the collision frequency between the self-propelled particle and the inner carrier wall, resulting in a larger effective active pressure and a more pronounced impact on pore opening.

Third, our results inform on the mechanisms and thermodynamics of pore formation in vesicles of distinct sizes and identify routes to potentially accelerate the process. We show that the tensionless exit barrier grows with the particle size, from ≈ 68*k*_*B*_*T* for MP_60_ to ≈ 110*k*_*B*_*T* for MP_180_, in line with previous results of swelling nanoparticles of about the same size^8^. Additionally, the larger nanoparticle remains trapped in a metastable pore-forming conformation, which does not widen to allow release because the line-tension cost of the rim grows faster than the curvature relief from opening^10^. In the *inside-out* active system, the confined self-propelled particle acts as a localized nonequilibrium force, as for active particles under confinement it pushes against the enclosing wall^61–63^, which propagates though the nanocarrier shell to the vesicle membrane, inducing pore widening and ultimately release for the smaller nanocarrier. The fully active case is less surprising mechanistically, as it more closely resembles a global external driving that acts over the entire carrier surface, analogous in spirit to field-driven membrane permeabilization. This distinction is relevant for nanomedical applications, including externally controlled release and cancer therapy, where localized forcing or electroporation-like mechanisms can be exploited to enhance membrane disruption and intracellular delivery^64–66^.

In conclusion, our results inform the physical mechanisms and thermodynamics of pore formation in vesicles of distinct sizes and identify routes to accelerate the process, aiding in the design of nanoparticles for enhanced release. The emerging picture is that release is governed by three coupled elements, the nature of the particle-membrane interaction, the size of the encapsulated particle and the spatial organization of the active driving force. A small nanoparticle (MP_60_) can be released passively once the repulsion is strong enough to destabilize the surrounding vesicle and open a pore wider than the particle radius, whereas the larger nanoparticle remains trapped in a metastable state even when a pore forms. Activity can lower the barrier and promote earlier pore opening, but its effectiveness depends on the application mode, *i*.*e*., a localized internal source assists release of the smaller particle, while only activity distributed over the whole nanoparticle sustains the pore expansion required to release the larger one. These results provide physical design principles for engineering nanocarriers with programmable, size-selective release.

## V. ACKNOWLEDGMENTS

I.M.I. acknowledges support from the Sectorplan Bèta & Techniek of the Dutch Government, the Dementia Research - Synapsis Foundation Switzerland and the Molecular Material Design Technology Hub. We thank Dr. Wouter den Otter for useful discussions.

## VI. DATA AVAILABILITY

The input files are publicly available on: https://github.com/ilieim/NanoparticleRelease

## AUTHOR CONTRIBUTIONS

M.P.: Conceptualization, Methodology, Software, Formal analysis, Investigation, Data curation, Visualization, Writing – review & editing.

I.M.I.: Conceptualization, Methodology, Supervision,Investigation, Project administration, Writing – review & editing, Funding acquisition.

## References

1 M. Deserno, Phys. Rev. E 69 (2004).

2 E. J. Spangler, S. Upreti, and M. Laradji, J. Chem. Phys. 144 (2016).

3 A. H. Bahrami, M. Raatz, J. Agudo-Canalejo, R. Michel, E. M. Curtis, C. K. Hall, M. Gradzielski, R. Lipowsky, and T. R. Weikl, Adv. Col. and Interface Sci. 208, 214–224 (2014).

4 S. A. Smith, L. I. Selby, A. P. R. Johnston, and G. K. Such, Bioconjugate Chemistry 30, 263–272 (2018).

5 X. A. Wu, C. H. J. Choi, C. Zhang, L. Hao, and C. A. Mirkin, Journal of the American Chemical Society 136, 7726–7733 (2014).

6 S. Zhang, H. Gao, and G. Bao, ACS Nano 9, 8655 (2015).

7 T. Debnath, J. Midya, T. Auth, and G. Gompper, ACS Macro Letters, 1412–1417 (2025).

8 R. Vácha, F. J. Martinez-Veracoechea, and D. Frenkel, ACS Nano 6, 10598–10605 (2012).

9 T. V. Tolpekina, W. K. den Otter, and W. J. Briels, J. Chem. Phys. 121, 8014–8020 (2004).

10 T. V. Tolpekina, W. K. den Otter, and W. J. Briels, J. Chem. Phys. 121, 12060–12066 (2004).

11 L. J. Starke, C. Allolio, and J. S. Hub, PNAS Nexus 4 (2025).

12 M. Post and G. Hummer, Nature Communications 17 (2025).

13 D. T. Zhang, L. Baldauf, G. Lazarski, T. S. van Erp, and W. Shinoda, J. Chem. Theor. Comp. 22, 3072–3083 (2026).

14 W. Shinoda, T. Nakamura, and S. O. Nielsen, Soft Matter 7, 9012 (2011).

15 A. Verma, O. Uzun, Y. Hu, Y. Hu, H.-S. Han, N. Watson, S. Chen, J. Irvine, and F. Stellacci, Nature Materials 7, 588–595 (2008).

16 D. Pei, ACS Nano 19, 40293–40303 (2025).

17 C. Qiu, F. Xia, J. Zhang, Q. Shi, Y. Meng, C. Wang, H. Pang, L. Gu, C. Xu, Q. Guo, and J. Wang, Research 6 (2023).

18 Y. Wang, V. Ukwattage, Y. Xiong, and G. K. Such, Materials Horizons 12, 3622–3632 (2025).

19 Z. Li, S. Tan, S. Li, Q. Shen, and K. Wang, Oncology Reports 38, 611–624 (2017).

20 K. Bhattacharjee and B. L. V. Prasad, Chem. Soc. Rev. 52, 2573–2595 (2023).

21 Y. Kumar, A. S. K. Sinha, K. D. P. Nigam, D. Dwivedi, and J. S. Sangwai, Nanoscale 15, 6075–6104 (2023).

22 I. Hafez, N. Maurer, and P. Cullis, Gene Therapy 8, 1188–1196 (2001).

23 S. C. Semple, A. Akinc, J. Chen, A. P. Sandhu, B. L. Mui, C. K. Cho, D. W. Y. Sah, D. Stebbing, E. J. Crosley, E. Yaworski, I. M. Hafez, J. R. Dorkin, J. Qin, K. Lam, K. G. Rajeev, K. F. Wong, L. B. Jeffs, L. Nechev, M. L. Eisenhardt, M. Jayaraman, M. Kazem, M. A. Maier, M. Srinivasulu, M. J. Weinstein, Q. Chen, R. Alvarez, S. A. Barros, S. De, S. K. Klimuk, T. Borland, V. Kosovrasti, W. L. Cantley, Y. K. Tam, M. Manoharan, M. A. Ciufolini, M. A. Tracy, A. de Fougerolles, I. MacLachlan, P. R. Cullis, T. D. Madden, and M. J. Hope, Nature Biotechnology 28, 172–176 (2010).

24 O. Boussif, F. Lezoualc’h, M. A. Zanta, M. D. Mergny, D. Scherman, B. Demeneix, and J. P. Behr, Proceedings of the National Academy of Sciences 92, 7297–7301 (1995).

25 N. Sonawane, F. C. Szoka, and A. Verkman, Journal of Biological Chemistry 278, 44826–44831 (2003).

26 L. Yang, T. A. Harroun, T. M. Weiss, L. Ding, and H. W. Huang, Biophys. J. 81, 1475–1485 (2001).

27 G. Tesei, Y.-W. Hsiao, A. Dabkowska, G. Grönberg, M. Yanez Arteta, D. Ulkoski, D. J. Bray, M. Trulsson, J. Ulander, M. Lund, and L. Lindfors, Proc. Nat. Acad. Sci. 121 (2024).

28 J. M. Johansson, H. Du Rietz, H. Hedlund, H. C. Eriksson, E. Oude Blenke, A. Pote, S. Harun, P. Nordenfelt, L. Lindfors, and A. Wittrup, Nature Communications 16 (2025).

29 B. Leonardini, M. Accorsi, R. Dimova, A. Relini, G. Rossi, and E. Canepa, Advances in Physics: X 11 (2026).

30 L. I. Selby, C. M. Cortez-Jugo, G. K. Such, and A. P. Johnston, WIREs Nanomedicine and Nanobiotechnology 9 (2017).

31 E. Sulheim, H. Baghirov, E. von Haartman, A. Bøe, A. K. O. Åslund, Y. Mørch, and C. d. L. Davies, J. Nanobiotech. 14 (2016).

32 L. Baldauf, F. Frey, M. Arribas Perez, T. Idema, and G. H. Koenderink, Biophysical Journal 122, 2311–2324 (2023).

33 J. Gilleron, W. Querbes, A. Zeigerer, A. Borodovsky, G. Marsico, U. Schubert, K. Manygoats, S. Seifert, C. Andree, M. Stöter, H. Epstein-Barash, L. Zhang, V. Koteliansky, K. Fitzgerald, E. Fava, M. Bickle, Y. Kalaidzidis, A. Akinc, M. Maier, and M. Zerial, Nature Biotechnology 31, 638–646 (2013).

34 G. Sahay, W. Querbes, C. Alabi, A. Eltoukhy, S. Sarkar, C. Zurenko, E. Karagiannis, K. Love, D. Chen, R. Zoncu, Y. Buganim, A. Schroeder, R. Langer, and D. G. Anderson, Nature Biotechnology 31, 653–658 (2013).

35 P. Gao, X. Jiang, J. Li, J. Nicolas, and T. Ha-Duong, Advanced Healthcare Materials 15 (2025).

36 S. Izvekov, A. Violi, and G. A. Voth, J. Phys. Chem. B 109, 17019–17024 (2005).

37 S. W. Park, B. H. Lee, and M. K. Kim, Mult. Sci. and Eng. 5, 104–118 (2023).

38 S. Franco-Ulloa, D. Guarnieri, L. Riccardi, P. P. Pompa, and M. De Vivo, J. Chem. Theor. Comp. 17, 4512–4523 (2021).

39 V. Schubertová, F. J. Martinez-Veracoechea, and R. Vácha, Soft Matter 11, 2726–2730 (2015).

40 F. Simonelli, D. Bochicchio, R. Ferrando, and G. Rossi, J. Phys. Chem. Lett 6, 3175–3179 (2015).

41 J. Lin, L. Miao, G. Zhong, C.-H. Lin, R. Dargazangy, and A. Alexander-Katz, Commun. Biol. 3.

42 S. Dixit, F. Noé, and T. R. Weikl, eLife 14 (2025).

43 I. R. Cooke, K. Kremer, and M. Deserno, Phys. Rev. E 72, 011506 (2005).

44 I. R. Cooke and M. Deserno, Biophys. J. 91, 487–495 (2006).

45 I. R. Cooke and M. Deserno, J. Chem. Phys. 123 (2005).

46 M. Paesani and I. M. Ilie, J. Chem. Phys. 164 (2026).

47 R. Vácha, F. J. Martinez-Veracoechea, and D. Frenkel, Nanolett. 11, 5391–5395 (2011).

48 B. J. Reynwar, G. Illya, V. A. Harmandaris, M. M. Müller, K. Kremer, and M. Deserno, Nature 447, 461–464 (2007).

49 S. Salassi, L. Caselli, J. Cardellini, E. Lavagna, C. Montis, D. Berti, and G. Rossi, J. Chem. Theor. Comp. 17, 6597–6609 (2021).

50 G. Rossi, P. F. J. Fuchs, J. Barnoud, and L. Monticelli, The Journal of Physical Chemistry B 116, 14353–14362 (2012).

51 T. R. Weikl, Soft Matter 22, 4424–4432 (2026).

52 M. Paesani and I. M. Ilie, J. Chem. Phys. 161, 244905 (2024).

53 M. Grau and E. Wagner, Current Opinion in Chemical Biology 81, 102506 (2024).

54 S. Rayamajhi, J. Marchitto, T. D. T. Nguyen, R. Marasini, C. Celia, and S. Aryal, Colloids and Surfaces B: Biointerfaces 188, 110804 (2020).

55 N. Ponomareva, S. Brezgin, I. Karandashov, A. Kostyusheva, P. Demina, O. Slatinskaya, E. Bayurova, D. Silachev, V. S. Pokrovsky, V. Gegechkori, E. Khaydukov, G. Maksimov, A. Frolova, I. Gordeychuk, A. A. Zamyatnin Jr., V. Chulanov, A. Parodi, and D. Kostyushev, Pharmaceutics 16, 667 (2024).

56 J. Litster, Physics Letters A 53, 193–194 (1975).

57 W. F. D. Bennett, N. Sapay, and D. P. Tieleman, Biophys. J. 106, 210–219 (2014).

58 M. Hu, J. J. Briguglio, and M. Deserno, Biophys. J. 102, 1403–1410 (2012).

59 P.-A. Boucher, B. Joós, M. J. Zuckermann, and L. Fournier, Biophys. J. 92, 4344–4355 (2007).

60 Y. Roiter, M. Ornatska, A. R. Rammohan, J. Balakrishnan, D. R. Heine, and S. Minko, Nano Letters 8, 941–944 (2008).

61 S. Das, G. Gompper, and R. G. Winkler, New Journal of Physics 20, 015001 (2018).

62 R. G. Winkler, A. Wysocki, and G. Gompper, Soft Matter 11, 6680–6691 (2015).

63 T. F. Zhu and J. W. Szostak, Journal of Systems Chemistry 2 (2011).

64 M. P. Stewart, R. Langer, and K. F. Jensen, Chem. Rev. 118, 7409–7531 (2018).

65 A. Rodzinski, R. Guduru, P. Liang, A. Hadjikhani, T. Stewart, E. Stimphil, C. Runowicz, R. Cote, N. Altman, R. Datar, and S. Khizroev, Sci. Rep. 6 (2016).

66 X. Zhang and A. G. Ewing, ACS Nano 16, 9852–9858 (2022).

