## Supplementary Information for "Physical mechanisms of nanoparticle intracellular release through membrane pore nucleation and expansion"

### I. SIMULATION DETAILS

To investigate the intracellular release of a nanoparticle from a vesicle, we use our recently introduced *metaparticle* model for flexible nanocarriers[1] combined with the Cooke-Deserno three-bead lipid model[2]. The initial conformation is selected from a successful endocytosis event from our previous work[3]. In the metaparticle model, a nanocarrier is represented as a collection of beads connected by spring-like bonds and arranged in highly symmetric topologies. Here, we use two models to represent nanoparticles of identical icosahedral topology but of distinct sizes, reflected in the number of beads in their composition, *i.e.,* 60 and 180, for $MP_{60}$ and $MP_{180}$, respectively (Fig. S1). The beads are interconnected by FENE springs with a spring constant $K = 30\,\varepsilon/\sigma^2$ and a maximum bond extension 1.8 $\sigma$ and sterically repel each other according to a short-range repulsive Weeks-Chandler-Andersen potential[1]. The two topologies correspond to effective radii of 4.5 $\sigma$ ($MP_{60}$) and 7 $\sigma$ ($MP_{180}$), with $\sigma$ the unit of length, here taken as 1 nm. Non-bonded beads interact via a truncated Lennard-Jones potential, with an effective bead diameter $\bar{\sigma} = 2.5\sigma$, a cut-off distance of 2.5 $\sigma$ and an interaction strength $\varepsilon_{\mathrm{LJ}} = 1\ k_BT$.

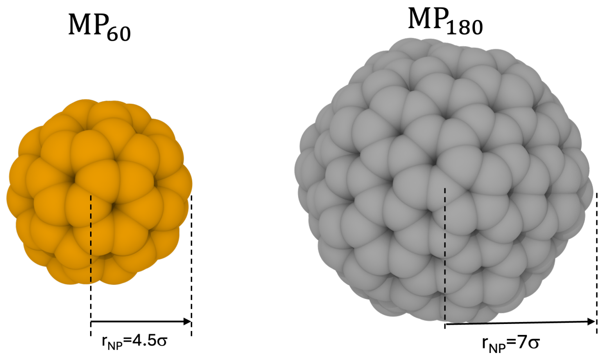


Fig. S1. **Nanoparticle models considered in this work.** Representative configurations of $MP_{60}$ and $MP_{180}$, composed of 60 and 180 interconnected beads, respectively. The two nanocarriers share the same icosahedral topology but differ in size, with effective radii of approximately 4.5 $\sigma$ for $MP_{60}$ and 7.0 $\sigma$ for $MP_{180}$.

The nanocarrier interacts with a lipid membrane, modeled using the coarse-grained lipid model introduced by Cooke and Deserno[2], which we and others have previously used to investigate key membrane-mediated processes such as wrapping, fusion and direct translocation[3–5]. In the Cooke-Deserno framework, each lipid is represented by three beads, *i.e.,* one hydrophilic bead corresponding to the head group and two hydrophobic beads corresponding to the tails. Bonded neighbors within a lipid are held together by a FENE potential with spring constant 30 $\varepsilon/\sigma^2$ and a maximum extension $r_0 = 1.5\sigma$. together with a short-range WCA repulsion, while a harmonic bending potential of stiffness $k_\mathrm{a} = 10\varepsilon$ acting between the head bead and the terminal tail bead controls the lipid stiffness. Bilayer cohesion and fluidity arise from an attractive cosine potential of range $w_c = 1.6\sigma$ acting between tail beads, a choice known to yield a stable fluid membrane[2]. The effective interaction diameters are $\bar{\sigma} = 0.95\sigma$ for head-head and head–tail pairs and $\bar{\sigma} = \sigma$ for tail-tail pairs. At these parameters the model self-assembles into a fluid bilayer of thickness $t = 5\sigma$. For the parametrisation used here, the bending rigidity, the Gaussian modulus and the edge line tension have been measured directly as $\kappa = 13k_BT$, $\kappa_G = -11.5k_BT$ and $\gamma = 3k_BT/\sigma$, respectively[6].

The interaction between a metaparticle and the membrane is described by a modified Lennard-Jones potential:

$$\Phi_{\mathrm{NB}}^{\mathrm{MP\text{-}mbr}}(r) = \mu(r,\bar{\sigma})\left[\varepsilon_{\mathrm{rep}}\left(\frac{\bar{\sigma}}{r}\right)^{12} - \varepsilon_{\mathrm{attr}}\left(\frac{\bar{\sigma}}{r}\right)^{6}\right] \quad \text{(S1)}$$

with the effective interaction diameter between each MP bead and membrane bead determined using the Lorentz-Berthelot mixing rules[7], $\varepsilon_{\mathrm{rep}}$ the repulsive interaction strength and $\varepsilon_{\mathrm{attr}}$ the attractive component of the potential. $\mu(r,\bar{\sigma}) = \mu(r,\bar{\sigma}) = \frac{1}{2}\left[1 - \frac{\tanh\left(\alpha(r-2\bar{\sigma})\right)}{\tanh\left(2\alpha\bar{\sigma}\right)}\right]$ is a step function that smoothly decays from 1 to 0 and was introduced to ensure continuity of the potential and the forces, and $\alpha$ controls the sharpness of the transition region, set to $\alpha = 5$. Throughout this work, $\varepsilon_{\mathrm{attr}} = 1\varepsilon$ toward both the lipid head and tail beads, the repulsion towards the lipid tail beads is set to $\varepsilon_{\mathrm{rep}}^{\mathrm{T}} = 2\varepsilon$ and the repulsion towards the lipid heads $\varepsilon_{\mathrm{rep}}^{\mathrm{H}}$ is varied in the 1-9 $\varepsilon$ interval. For ease of representation we use $\varepsilon_{\mathrm{rep}} = \varepsilon_{\mathrm{rep}}^{\mathrm{H}}$ in the main text (see Fig. S2).

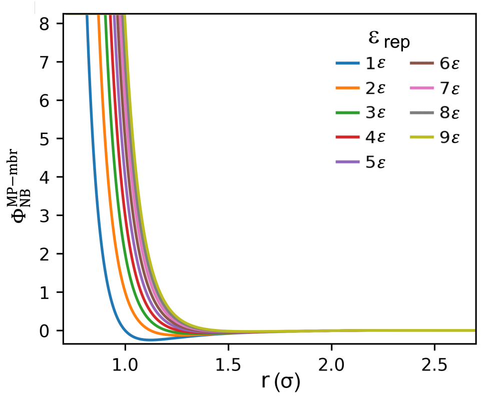


Fig. S2. **MP-lipid head beads interaction potential.** Shown are profiles for $\varepsilon_{\mathrm{rep}} \in 1\varepsilon$ to $9\varepsilon$ at constant $\varepsilon_{\mathrm{attr}} = 1\varepsilon$ and a cut-off distance $r_c = 4.1\ \sigma$.

#### II. PORE RADII

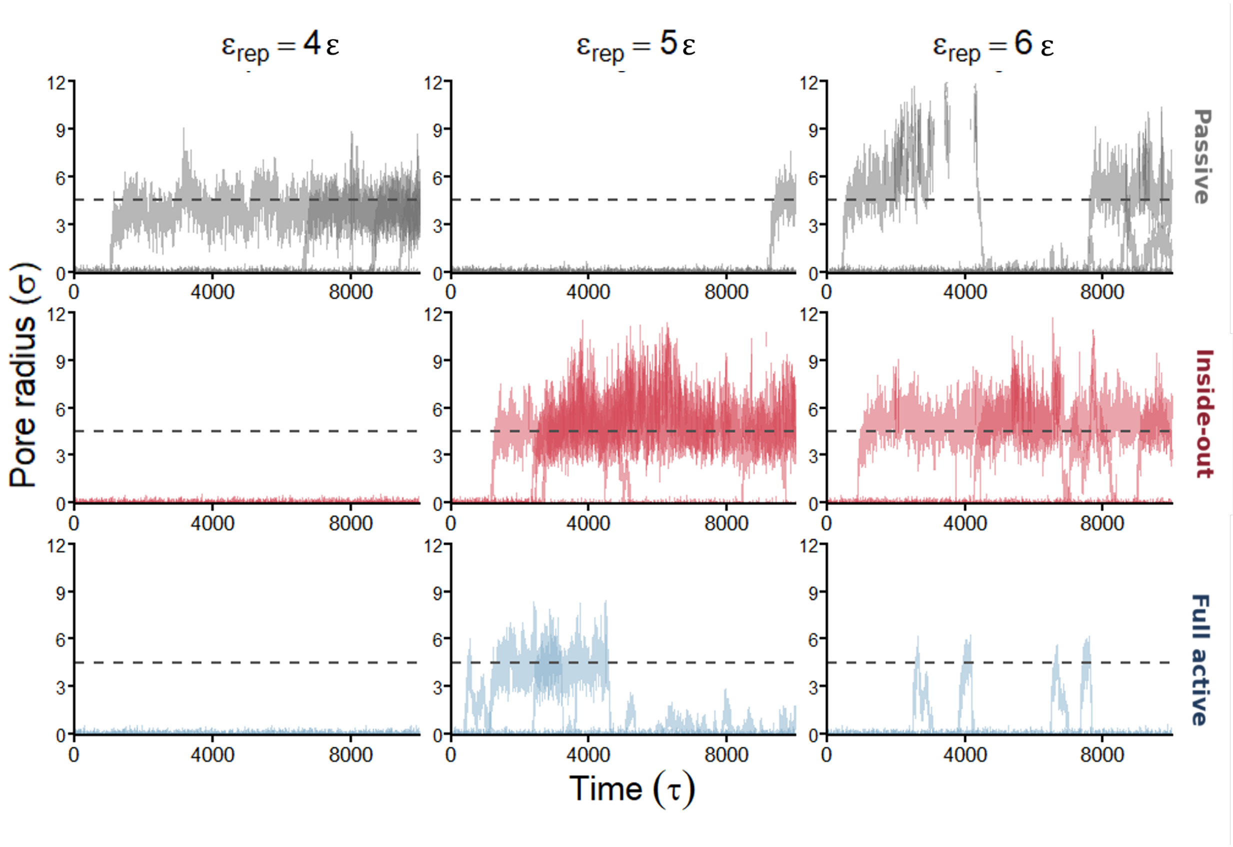


Fig. S3. **Time evolution of the effective pore radius for $MP_{60}$.** The columns correspond to $\varepsilon_{rep} = 4\varepsilon$, $5\varepsilon$, and $6\varepsilon$, while the rows show the passive (grey prfiles), inside-out active (red profiles) and fully active release mechanisms (blue profiles), respectively. Each curve represents one of five independent simulations. The horizontal dashed line indicates the reference nanoparticle radius of $4.5\sigma$.

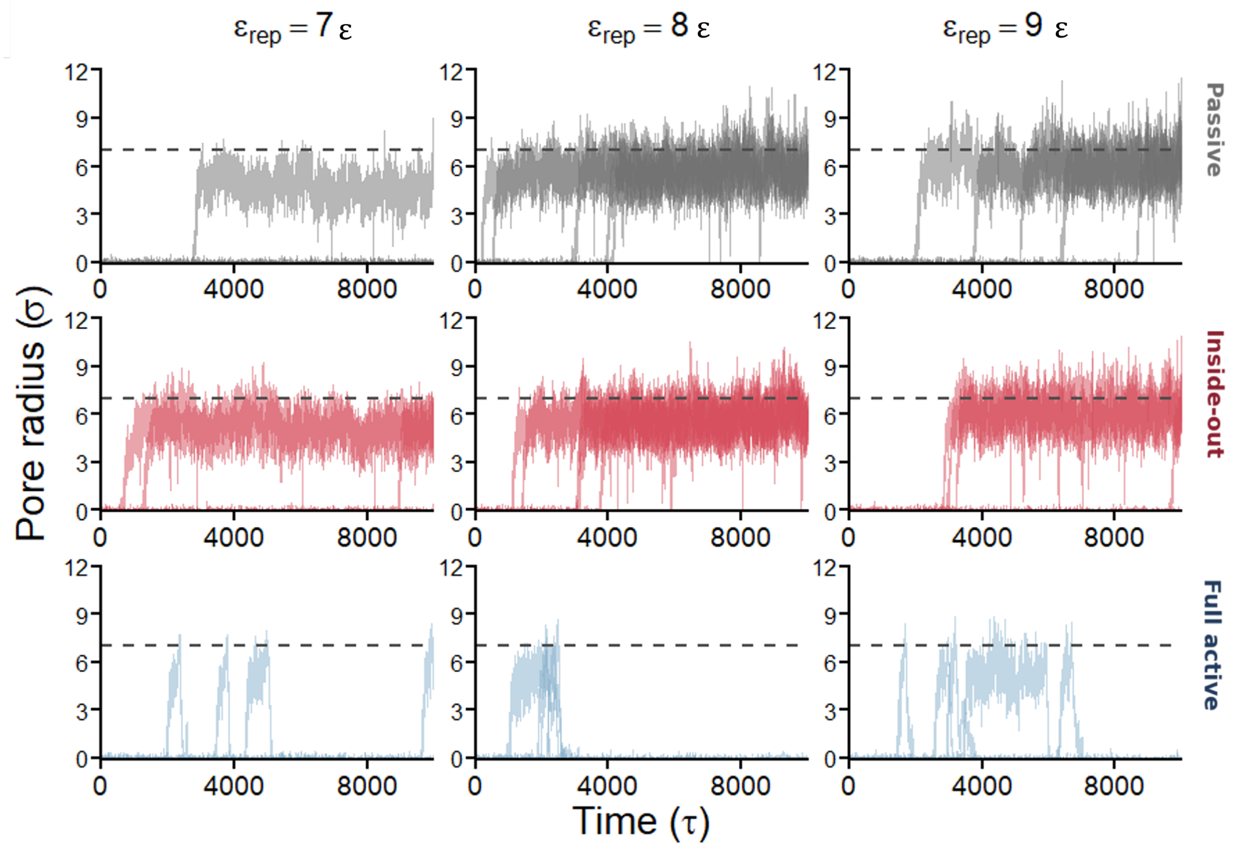


Fig. S4. **Time evolution of the effective pore radius for $MP_{180}$ at high MP-membrane repulsion.** The columns correspond to $\varepsilon_{rep} = 7\varepsilon$, $8\varepsilon$, and $9\varepsilon$, while the rows show the passive (grey profiles), inside-out active (red profiles) and fully active release mechanisms (blue profiles), respectively. Each curve represents one of five independent simulations. The horizontal dashed line marks the $MP_{180}$ radius, $r_{NP} = 7\sigma$.

#### 53 III. ROLLING EXIT STRATEGY

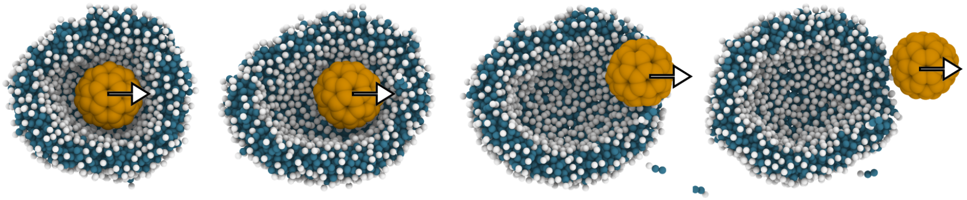

Fig. S5. **Rolling escape of a fully active nanoparticle from the surrounding vesicle.** Representative sequence showing the release of $MP_{60}$ under full activation. The active force, homogeneously applied to all metaparticle beads, drives the nanocarrier along the direction indicated by the arrows. The resulting membrane deformation is not localized along the nanocarrier displacement axis, and pore opening occurs perpendicular to the view plane and the activity direction, allowing the nanoparticle to roll along the pore edge and escape from the vesicle.

#### IV. PORE FREE-ENERGY CALCULATION

We estimate the free-energy cost of pore formation in the passive vesicle using the classical continuum description of a circular pore in an elastic membrane[8–11].

##### A. Parameters

All quantities are expressed in the simulation units of length $\sigma$ and energy $k_BT$.

| Quantity | Symbol | Value | Reference |
|---|---|---|---|
| Bending rigidity | $\kappa$ | $13k_BT$ | Ref.[6] |
| Gaussian rigidity | $\kappa_G$ | $-11.5k_BT$ | Ref.[6] |
| Pore line tension | $\gamma$ | $3k_BT/\sigma$ | Ref.[6] |
| Area-stretch modulus | $K_S$ | $250\mathrm{mJ\,m^{-2}} = 60.4\ k_BT/\sigma^2$ | Ref.[9] |
| Membrane thickness | $t$ | $5\sigma$ | Refs.[2,3] |
| Vesicle midplane radius ($\mathrm{MP}_{60}$) | $R_0$ | $7.4\sigma$ | this work |
| Vesicle midplane radius ($\mathrm{MP}_{180}$) | $R_0$ | $10.05\sigma$ | this work |
| Particle radius ($\mathrm{MP}_{60}$) | $r_{\mathrm{NP}}$ | $4.5\sigma$ | this work |
| Particle radius ($\mathrm{MP}_{180}$) | $r_{\mathrm{NP}}$ | $7.0\sigma$ | this work |

The calculated area-stretch modulus $K_S = 250\ \mathrm{mJ\,m^{-2}}$[9] is consistent with the $\sim 200\ \mathrm{mJ\,m^{-2}}$ obtained for phospholipid bilayers by micropipette aspiration[12,13]. The midplane radius is obtained from the measured outer radius as $R_0 = R_{\mathrm{out}} - t/2$.

###### Pore free energy on a closed tensionless vesicle

To open a pore of radius $r_p$ in a closed, tensionless vesicle, the free energy difference between the closed vesicle with radius $R_0$ and an opened vesicle with radius $R$ is[11]:

$$\Delta G(r_p) = 2\pi\gamma r_p - (2\kappa + \kappa_G)\,\frac{\pi r_p^2}{R_0^2}. \tag{S2}$$

Releasing the particle requires expanding the pore to the particle size, $r_p = r_{\mathrm{NP}}$. Hence, the barrier maximum height would correspond to a critical radius

$$r^* = \frac{\gamma R_0^2}{2\kappa + \kappa_G}. \tag{S3}$$

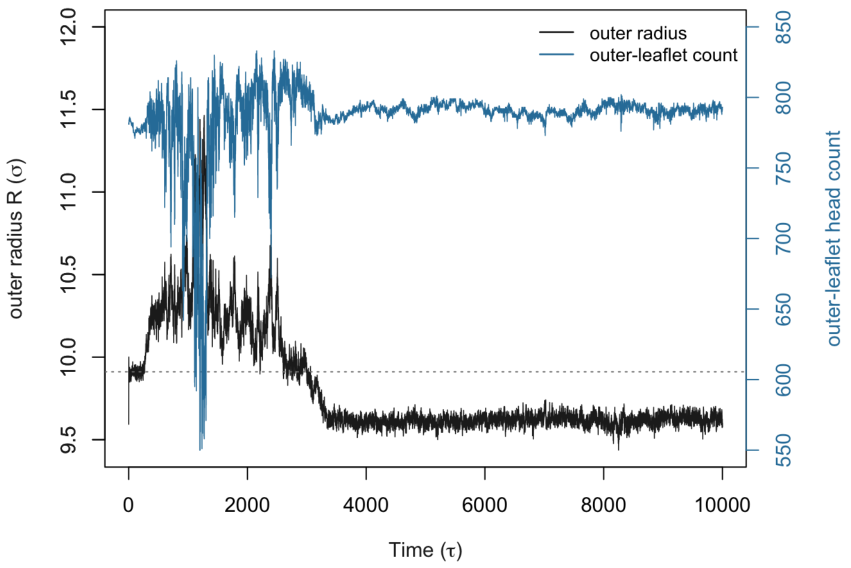


Fig. S6. Timeline of the outer vesicle radius (black line and y-axis on the left) and outer-leaflet head-bead count (blue line and y-axis on the right) for passive $MP_{60}$ at $\varepsilon_{rep} = 7\varepsilon$, for which the nanoparticle is released from the enclosing vesicle.

For the nanoparticles studied here, $r^*$ exceeds the vesicle radius to $r^* \approx 11.3\sigma$ for $MP_{60}$ ($R_0 = 7.4\sigma$) and $r^* \approx 20.7\ \sigma$ for $MP_{180}$ ($R_0 = 10\sigma$).

##### Nucleation barrier under transient stretch

In the early stage of the $\varepsilon_{rep} = 7\varepsilon$ trajectory the $MP_{60}$ vesicle swells while remaining closed (Fig. S6). Essentially the outer-leaflet radius grows but the outer-leaflet lipid count stays near its baseline value of about 780 head-beads (Fig. 4(b)). We quantify this stretch through the area strain of the outer leaflet, defined from the instantaneous outer-leaflet radius $R_i$ and lipid count $N_i$, as the fractional change in area per lipid relative to the wrapped (unstretched) reference state $(R_0, N_0)$,

$$\tilde{\varepsilon} = \frac{a_i}{a_0} - 1 = \frac{4\pi R_i^2/N_i}{4\pi R_0^2/N_0} - 1 = \frac{R_i^2}{R_0^2} \cdot \frac{N_0}{N_i} - 1, \tag{S4}$$

where $a_i = 4\pi R_i^2/N_i$ is the instantaneous area per outer-leaflet lipid and $a_0 = 4\pi R_0^2/N_0$ its value in the wrapped state ($R_{out} \approx 9.9\sigma$, $N_0 \approx 780$). During the closed-swollen window the radius increases to $R_i \approx 10.15\sigma$ at constant $N_i$, resulting in a transient area strain $\tilde{\varepsilon} \approx 5\%$.

This strain imposes a transient membrane tension

$$\Lambda = K_S\,\tilde{\varepsilon} \approx 3\ k_B T/\sigma^2. \tag{S5}$$

On a closed vesicle, the curvature relief accompanying pore opening adds a further effective tension, and the resulting total tension driving nucleation is $\Lambda_{\text{tot}} = \Lambda + (2\kappa + \kappa_G)/R_0^2 \approx 3.3\ k_B T/\sigma^2$, which reduces to the known Litster equation in case of flat membranes[8,9]. The resulting nucleation barrier and critical pore radius are

$$\Delta G^*_{\text{nuc}} = \frac{\pi\gamma^2}{\Lambda_{\text{tot}}} \approx 9\ k_B T, \qquad r^*_{\text{nuc}} = \frac{\gamma}{\Lambda_{\text{tot}}} \approx 0.9\ \sigma, \tag{S6}$$

placing the barrier only a few $k_B T$ above thermal energy, so that a pore nucleates readily once the bilayer is stretched.

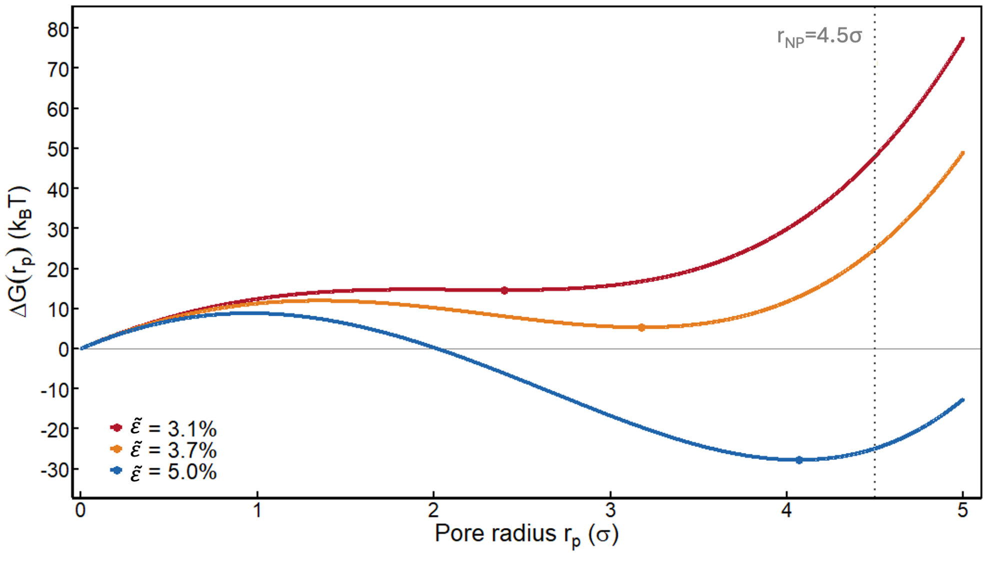


Fig. S7. The pore free energy $\Delta G(r_p)$ calculated from Eq. (S9) is shown as a function of the pore radius $r_p$ for outer-leaflet area strains $\tilde{\varepsilon} = 3.1\%$, 3.7%, and 5.0%. Highlighted are the local metastable minima of the free-energy profiles (filled circles), and the vertical dotted line indicates the $MP_{60}$ radius, $r_{\mathrm{NP}} = 4.5\,\sigma$, corresponding to the pore size required for escape.

#### Metastable pore free energy

Once a pore has nucleated, its subsequent evolution is governed by the elastic energy stored in the stretched outer leaflet, which the pore partially relaxes as it absorbs membrane area. We extend the work of Tolpekina *et al.*[10] for flat bilayers to account for curved vesicles. Essentially, we add to the line-tension and curvature terms a constant-area elastic term that penalises the residual area strain after a pore of area $\pi r_p^2$ has opened. We assume the bilayer carries no spontaneous curvature. As such, the neutral plane coincides with the bilayer midplane and the Helfrich energy density of the intact vesicle is $(2\kappa + \kappa_G)/R_0^2$. The free energy of the porated vesicle reads

$$G(r_p) = 2\pi\gamma r_p + (2\kappa + \kappa_G)\frac{A_0 - \pi r_p^2}{R_0^2} + \frac{K_S}{2A_0}\left(A_0\,\tilde{\varepsilon} - \pi r_p^2\right)^2, \tag{S7}$$

where $A_0 = 4\pi R_{\mathrm{out}}^2$ is the tensionless outer-leaflet area and $\tilde{\varepsilon}$ the area strain imposed on it (Eq. S4). The first two terms on the r.h.s. represent the contributions from the line tension and the curvature, respectively. The third term on the r.h.s. represents the elastic energy stored in the strained membrane, which is minimised when the pore has grown large enough to absorb the excess area, $\pi r_p^2 = A_0\tilde{\varepsilon}$.

The change in free energy relative to the intact pore-less vesicle is physically more meaningful, $\Delta G(r_p) = G(r_p) - G(0)$. Setting $r_p = 0$ in Eq. (S7) results in

$$G(0) = (2\kappa + \kappa_G)\,\frac{A_0}{R_0^2} + \frac{K_S}{2A_0}\,(A_0\,\tilde{\varepsilon})^2\,. \tag{S8}$$

Subtracting term by term, the change in free-energy is

$$\Delta G(r_p) = 2\pi\gamma r_p - \left[\frac{2\kappa + \kappa_G}{R_0^2} + K_S\,\tilde{\varepsilon}\right]\pi r_p^2 + \frac{K_S\,\pi^2}{2A_0}\,r_p^4. \tag{S9}$$

The competition between the linear line-tension term (opposing the opening) and the quadratic relief term (favouring the opening), re-stabilised by the quartic finite-reservoir term that penalises over-expansion once the excess area is absorbed, produces a metastable pore once the strain exceeds a threshold $\tilde{\varepsilon}^* \approx 3.1\%$ (Fig. S7). Specifically, at $\tilde{\varepsilon} \approx 3.1\%$ the minimum is at $r_p \approx 2.4\ \sigma$, which is $\approx 33\ k_BT$ below the exit condition at $r_p = r_{\mathrm{MP}}$ ($\Delta G[r_{\mathrm{NP}} = 4.5\ \sigma] - \Delta G[2.4\ \sigma] = 48 - 15 = 33\ k_BT$). At $\tilde{\varepsilon} \approx 3.7\%$ the minimum shifts to $r_p \approx 3.2\ \sigma$, and still lies $\approx 20\ k_BT$ below the exit condition. At $\tilde{\varepsilon} \approx 5\%$ the minimum is at $r_p \approx 4.1\ \sigma$, close to the nanoparticle radius $r_{\mathrm{NP}}$, and the remaining barrier collapses to a few $k_BT$.

[1] M. Paesani and I. M. Ilie, J. Chem. Phys. **161**, 244905 (2024).

[2] I. R. Cooke, K. Kremer, and M. Deserno, Phys. Rev. E **72**, 011506 (2005).

[3] M. Paesani and I. M. Ilie, J. Chem. Phys. **164** (2026).

[4] J. C. Shillcock and R. Lipowsky, Nat. Mater. **4**, 225 (2005).

[5] R. Vácha, F. J. Martinez-Veracoechea, and D. Frenkel, Nanolett. **11**, 5391–5395 (2011).

[6] M. Hu, J. J. Briguglio, and M. Deserno, Biophys. J. **102**, 1403–1410 (2012).

[7] H. A. Lorentz, Annalen der Physik **248**, 127–136 (1881).

[8] J. Litster, Physics Letters A **53**, 193–194 (1975).

[9] T. V. Tolpekina, W. K. den Otter, and W. J. Briels, J. Chem. Phys. **121**, 12060–12066 (2004).

[10] T. V. Tolpekina, W. K. den Otter, and W. J. Briels, J. Chem. Phys. **121**, 8014–8020 (2004).

[11] R. Vácha, F. J. Martinez-Veracoechea, and D. Frenkel, ACS Nano **6**, 10598–10605 (2012).

[12] E. Evans and D. Needham, J. Phys. Chem. **91**, 4219–4228 (1987).

[13] W. Rawicz, K. Olbrich, T. McIntosh, D. Needham, and E. Evans, Biophys. J. **79**, 328–339 (2000).
